# Reaction-Diffusion Model of Cortical Atrophy Spread during Early Stages of Alzheimer’s Disease

**DOI:** 10.1101/2020.11.02.362855

**Authors:** Sue Kulason, Michael I Miller, Alain Trouvé, Alzheimer’s Disease Neuroimaging Initiative

**Affiliations:** Center for Imaging Science, The Johns Hopkins University, Baltimore, MD, USA; Institute for Computational Medicine, The Johns Hopkins University, Baltimore, MD, USA; Department of Biomedical Engineering, The Johns Hopkins University, Baltimore, MD, USA; Kavli Neuroscience Discovery Institute, Johns Hopkins University, Baltimore, MD, USA; Centre Borelli, CNRS, Ecole Normale Supérieure Paris-Saclay, Université Paris-Saclay, Gif-sur-Yvette, France

## Abstract

1.

This study introduces a reaction-diffusion model of atrophy spread across the rhinal cortex during early stages of Alzheimer’s disease. Our finite elements model of atrophy spread is motivated by histological evidence of a spatio-temporally specific pattern of neurofibrillary tau accumulation, and evidence of grey matter atrophy correlating with sites of neurofibrillary tau accumulation. The goal is to estimate disease-related parameters such as the origin of atrophy, the speed at which atrophy spreads, and the stage of the disease. We solve a constrained optimization problem using the adjoint state method and gradient descent to match modeled cortical thickness to observed cortical thickness as calculated from 3T MRI scans. Simulation testing shows that disease-related parameters can be estimated accurately with as little as 2 years of annual observations, depending on the stage of the disease. Case studies of 3 subjects suggests that we can pinpoint the origin of atrophy to the anterior transentorhinal cortex, and that the speed of atrophy spread is less than 1 mm per year. In the future, this type of modeling could be useful to stage the progression of the disease prior to the onset of clinical symptoms.

**Author Summary:** Misfolded tau proteins are associated with Alzheimer’s disease. They are known to accumulate and spread across the rhinal cortex, which is an area of the temporal lobe. Recent imaging studies suggest that we can detect grey matter thinning that occurs in pattern similar to tau spread. In this study, we introduce a model of disease spread to examine where thinning begins, how fast it spreads, and the stage of the disease. The results show that the origin of thinning corresponds with the earliest known location of tau accumulation, and spreads at a rate of less than 1 mm per year. Future work may focus on staging the progression of the disease using this type of model.

## 3. Introduction

There is a consensus that Alzheimer’s disease (AD) begins years before the onset of clinical symptoms [19]. It is of interest to identify individuals at risk of developing AD and stage disease progression during the preclinical period, which is prior to the onset of clinical symptoms. Research efforts have been underway to develop robust, sensitive biomarkers that stage the progression and severity of the disease.

The field of computational anatomy has been working to refine shape measures and increase sensitivity to disease-related changes. While the exact mechanism is not understood, neurofibrillary tau tangles, particularly in the presence of beta amyloid plaques, are associated with decreased neuron counts and increased grey matter atrophy in AD [5]. Recently, spatially specific patterns of grey matter atrophy have been detected in structural MRI prior to the development of clinical symptoms [22, 14]. There is evidence that AD-related atrophy begins 8 to 10 years prior to symptom onset in the entorhinal cortex (ERC), and 2 to 4 years prior to symptom onset in the hippocampus [24]. Our recent work has further suggested that disease-related atrophy begins in the transentorhinal cortex (TEC) before the ERC [11]. This pattern of atrophy mirrors the accumulation of neurofibrillary tau tangles, which has been used to stage Alzheimer’s disease post-mortem [3]. In addition, a recent PET imaging study has shown tau tracer retention that similarly progressed in a Braak-stage like pattern beginning in the rhinal cortex [16].

It is hypothesized that neurofibrillary tau tangles propagate in a prionlike manner by cell release, diffusion, and cell uptake along axonal pathways [6, 4, 15, 17, 2, 1]. Molecular and cellular data suggest that tau tangles accumulate and seed the conversion of normal tau to misfold and aggregate [15, 7]. The structural and functional connectivity of the TEC may explain the pattern of neurofibrillary tau and atropy spread. The TEC is a major contributor of input into the anterior lateral ERC [12, 21], the anterior proximal subiculum bordering CA1 [12, 8], and the basolateral and baso-medial nuclei of the amygdala [13]. More locally, connectivity within the ERC has been studied in the context of grid cells, which are spatially organized. In particular, Layer II interneurons of the medial ERC have been shown to connect to and inhibit neighboring principal neurons [23], suggesting spatially-dependent interconnections in the rhinal cortex.

Given the spatially and temporally specific spread of neurofibrillary tau tangles that correlate with grey matter atrophy, we propose to model AD-related atrophy as a reaction-diffusion process across the rhinal cortex. We hypothesize that connectivity within the rhinal cortex can be modeled as a function of distance, and that the progression of AD can be modeled from longitudinal MRI data.

## 4. Results

### 4.1. Model of atrophy spread

The overarching goal of this model is to represent cortical thickness as a function of disease, as it spreads across the rhinal cortex. In order to accurately depict cortical thickness over time, we incorporate information about age-related atrophy and the variable, non-uniform thickness across this piece of cortex.

Let *s* represent a location in the spatial domain Ω, and *t* represent a time in the temporal domain [*t*_0_, *t*_1_]. We propose a model of cortical thickness atrophy such that cortical thickness *ρ*(*s, t*) has a sigmoidal relationship with the stage of the disease *b*(*s, t*), and aging λ_0_(*s*) *t*, as shown in Eqn 1. The disease stage *b*(*s,t*) ∈ (−∞, ∞) increases as the disease progressively gets worse. The rate of disease progression *∂_t_b*(*s, t*) is controlled by the disease intensity λ > 0 and local activity rate a(s,t), as shown in Eqn 2. Figure 1 shows the relationship between cortical thickness, disease stage, and local activity rate at a single location.

**Figure 1:**
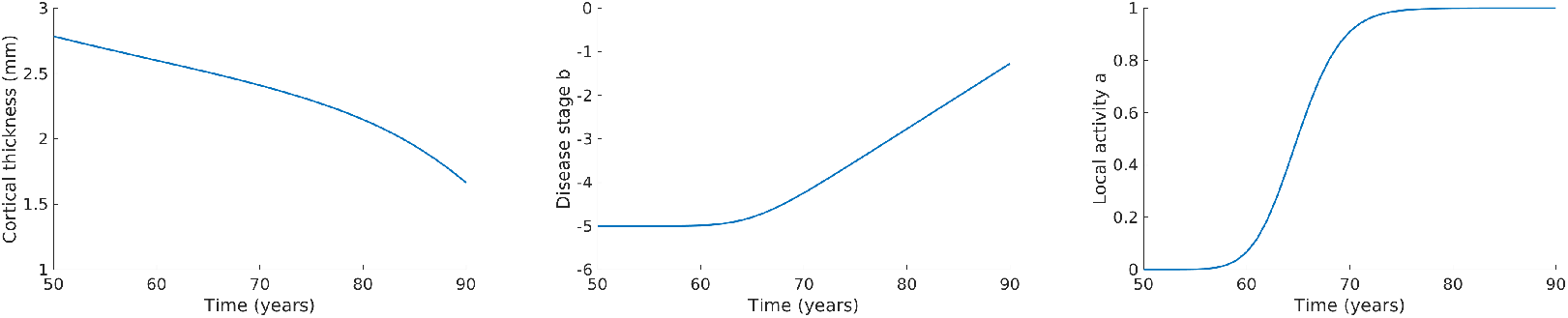
Cortical thickness (left), disease stage (middle), and local activity rate (right) for a point.

The local activity rate *a*(*s, t*) ∈ [0, 1] loosely represents the environmental conditions needed for disease progression, such as neurofibrillary tau tangle concentration. We model *a*(*s, t*) as a Fisher-KPP equation with a diffusion rate of *D* and reaction rate of *κ*, shown in Eqn 3. When *a*(*s, t*_0_) = 0 for all *s*, the disease will not progress. When *a*(*s*_0_, *t*_0_) is seeded with a non-zero value at some location *s*_0_, the model progresses to a stable fixed point of *a*(*s,t*_1_) = 1 for all s. At the boundaries of the manifold *s* ∈ *∂*Ω, we impose a Neumann boundary condition on the local activity rate *a*(*s,t*).

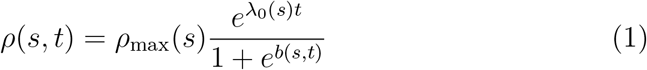

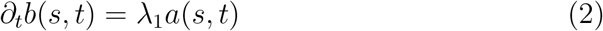

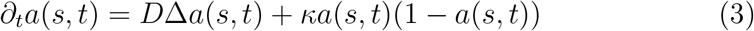

### 4.2. Disease simulation

We first determine age-related atrophy rate, sex differences, and the thickness of the rhinal cortex based on a set of stable NC subjects. Age-related atrophy rate had a mean of 0.75% per year and was as much as 1.62% per year in the posterior lateral region. Female subjects were up to 0.5 mm thicker than male subjects, and were thinner centrally. After removing the effects of age and sex, the cortex was thinnest in the posterior medial region, and thickest in the anterior central region. The average thickness at a given location was as much as 1. 46 mm away from the mean thickness across the cortex, strongly suggesting that cortical thickness cannot be accurately modeled as a uniform thickness across this region. Figure 2 shows the difference in thickness (shape difference), sex difference, and age-related atrophy rate. Details on data processing methods used to calculate cortical thickness are discussed in Section 6.1.

**Figure 2:**
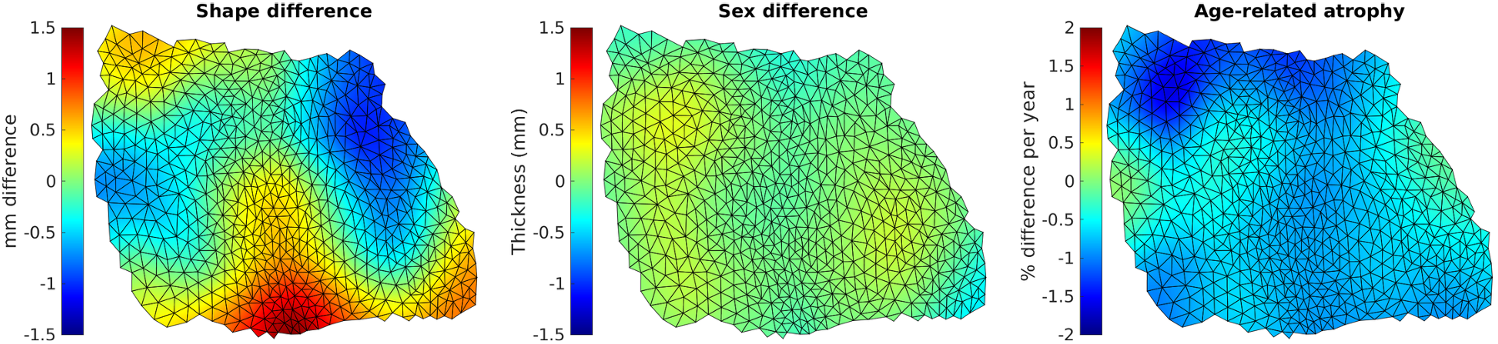
A priori model of shape difference (left), sex difference (middle) and age-related atrophy (right). Orientation of the surface is left (lateral), right (medial), top (posterior), bottom (anterior).

Next, we selected a set of disease parameters to simulate AD progression using the atrophy spread model (Eqn 1, 2, and 3). Parameter selection is discussed in Section 6.2. Figure 3 shows the model of age-related atrophy progression versus the simulation of AD progression. Qualitatively, there is a difference in thickness that grows noticeable decades after disease initiation at age 50.

**Figure 3:**
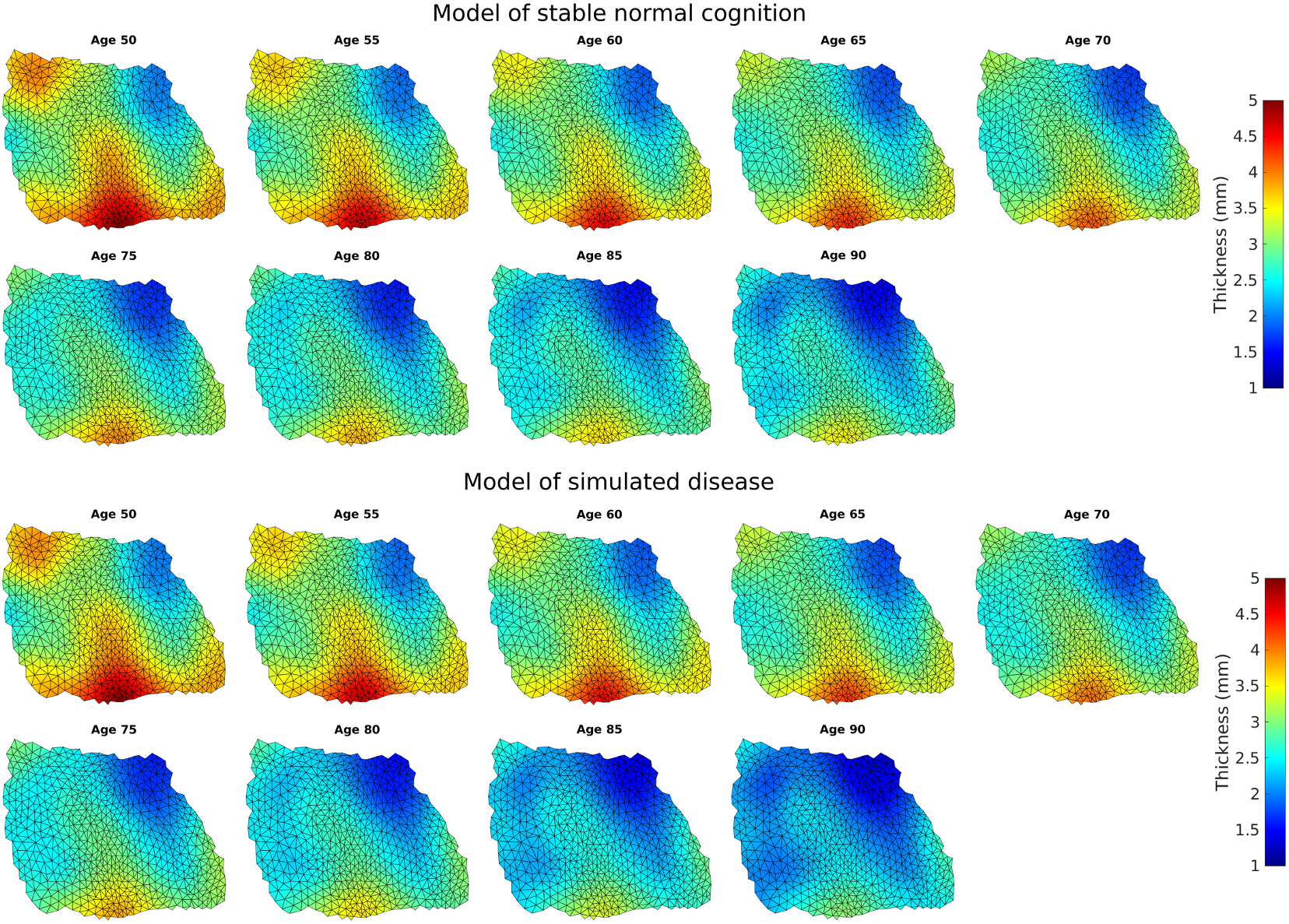
A priori model of cortical thickness for a female stable NC from age 50 to 90, top. Simulated disease for a female subject from age 50 to 90, bottom. Orientation of the surface is left (lateral), right (medial), top (posterior), bottom (anterior).

### 4.3. Parameter estimation on disease simulation

Here, we examine the effects of the window of observation on the accuracy of parameter estimation. The window of observation was selected to be a sliding window of annual observations between age 50 and 90, with a window size of 2, 5, and 10 years. For each window of observation on the disease simulation, we estimated model parameters from 100 random initial guesses. Figure 4 shows the fraction of runs where all parameters were estimated accurately.

**Figure 4:**
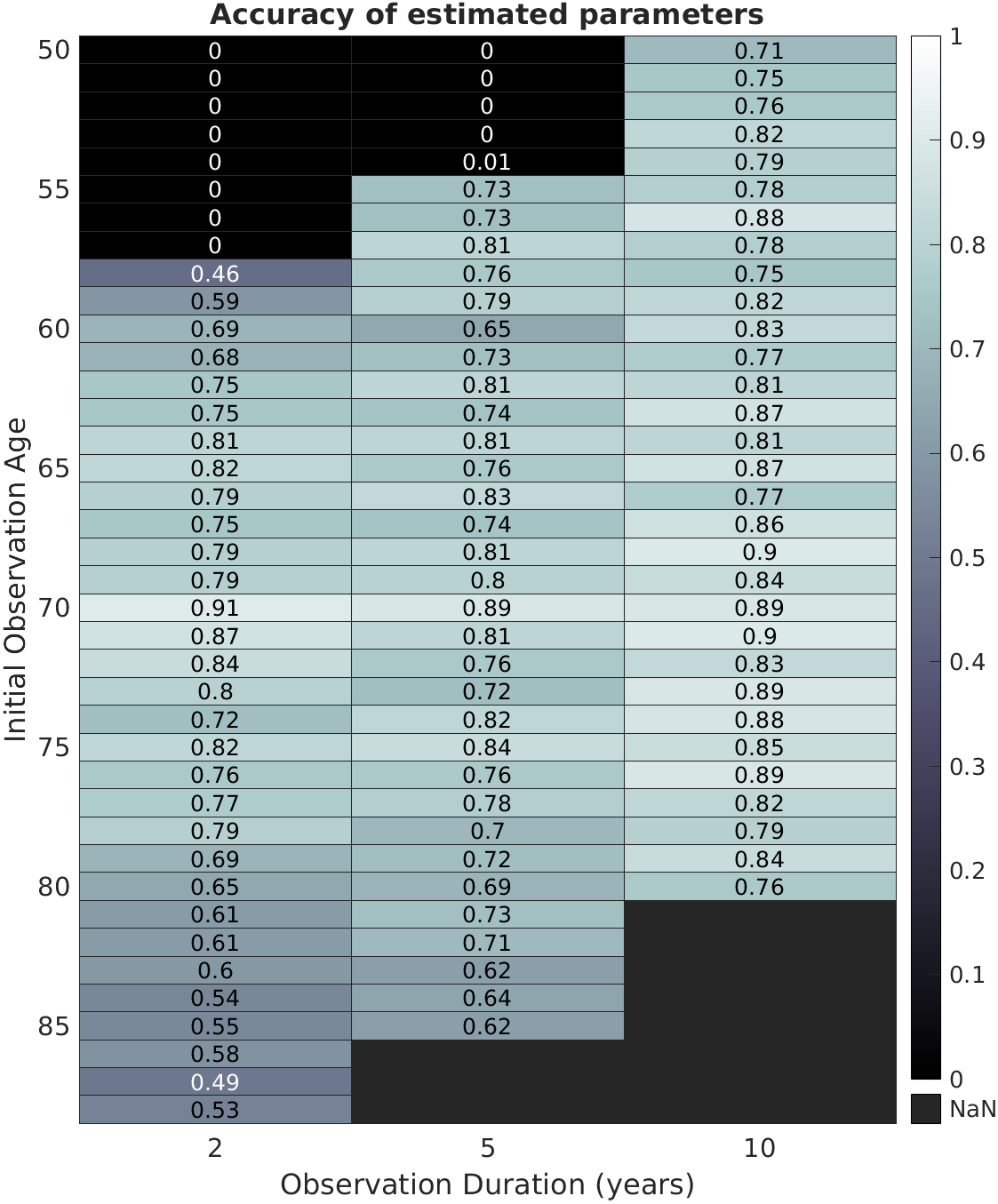
Fraction of 100 runs that succeeded in estimating the correct parameters over varying duration and observation start times

In general, greater accuracy was achieved with longer observation windows. The accuracy was poorest for windows of observation with an initial observation time before age 60. The accuracy was also poorer in windows of observation with an initial observation time after age 80. Best performance was achieved when observations started at age 70, 20 years after the initialization of this disease simulation. The maximum a posteriori (MAP) estimate for each window of observation estimated all parameters correctly, with a few exceptions. These were: observation windows of 2-year intervals from age 50 to 57, and 5-year intervals from age 50 to 53. This suggests that the parameters cannot be accurately estimated very early in the disease with a short window of observation.

In addition to examining the proportion of runs that accurately estimated all parameters, we assessed model accuracy by building confidence intervals for these parameters. A Metropolis-Hastings algorithm was implemented to approximate the posterior distribution, *p*(*θ* | *ρ*_observed_), starting from a set of observed cortical thicknesses, *ρ*_observed_. To derive an explicit expression for *p*(*θ* | *ρ*_observed_), we modeled noise in observations, *σ*^2^, as an inverse Wishart distribution of mean and variance 1. We then calculated the 90% confidence interval and the radius of a circle that circumscribed 90% of atrophy origins, *c*_0_. The details on this approach can be found in Section 6.4.

We compared the confidence intervals for three windows of observation: 1) early stage (annual observations from age 55 to 60), 2) mid stage (annual observations from age 70 to 75) and 3) late stage (annual observations from age 85 to 90). Table 1 shows the 90% confidence intervals that resulted from 20,000 samples.

**Table 1:**
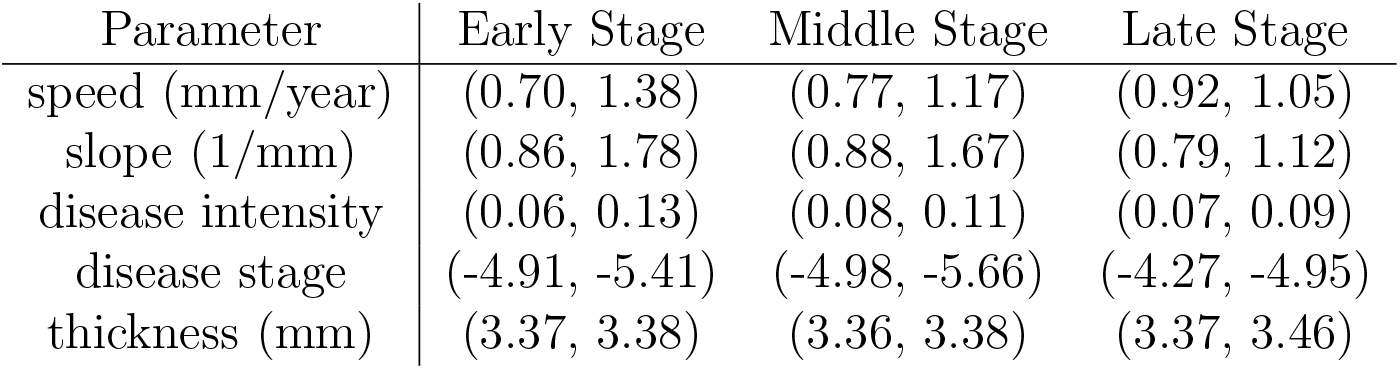
90% confidence interval for 5-year annual windows of observation in the early, middle and late stage of the disease.

| Parameter | Early Stage | Middle Stage | Late Stage |
| --- | --- | --- | --- |
| speed (mm/year) | (0.70, 1.38) | (0.77, 1.17) | (0.92, 1.05) |
| slope (1/mm) | (0.86, 1.78) | (0.88, 1.67) | (0.79, 1.12) |
| disease intensity | (0.06, 0.13) | (0.08, 0.11) | (0.07, 0.09) |
| disease stage | (-4.91, -5.41) | (-4.98, -5.66) | (-4.27, -4.95) |
| thickness (mm) | (3.37, 3.38) | (3.36, 3.38) | (3.37, 3.46) |

The radius that circumbscribed 90% of *c*_0_ was 0.48 mm for early stage, 0.58 mm for middle stage, and 0.36 mm for late stage. These are relatively small regions on the rhinal cortex. For context, the rhinal cortex is approximately 20 mm wide (medial to lateral) and 25 mm long (anterior to posterior).

Speed, 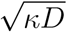, had the widest interval in the early stage before a substantial spread of atrophy could be observed, and the narrowest confidence interval in the late stage. The confidence interval for slope, 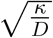, was also widest in the early stage and narrowest in the late stage. The confidence interval for speed is consistently narrower than for slope, suggesting slope is a more difficult parameter to estimate accurately.

Following the same trend as for speed and slope, disease intensity, λ_1_, had the widest confidence interval in the early stage and narrowest interval in the late stage. The confidence interval is narrow when the peak atrophy rate is captured during the observation window.

Initial disease stage, 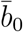, had the narrowest confidence interval in the early stage and widest confidence interval in the late stage. The initial disease stage is more accurately estimated when the window of observation is closer to initial age, which is age 50. Similarly, average cortical thickness prior to disease-related atrophy, 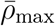, had the smallest confidence interval in the early stage and the largest confidence interval in late stage. The confidence interval is wider the further the observation window is from the initial age.

The results suggest that accurate estimations of the origin of atrophy, speed, and disease intensity can be established from 5 years of annual observations. Speed has somewhat poorer accuracy early in the disease, and initial disease stage and initial cortical thickness have somewhat poorer accuracy late in the disease.

### 4.4. Case study on real data

We now examine the cortical thickness of three preclinical AD subjects and fit them to the model of atrophy spread. Details on the selection and demographic details of these subjects can be found in Section 6.1. Figure 5 shows the observed cortical thicknesses, *ρ*_observed_, for each subject. Since the patterns of atrophy are difficult to see from scan to scan, we also show the change in cortical thickness with respect to the first scan in Figure 6. Subject 1 has a thinner cortex over all, but still shows the characteristic pattern of a thinner posterior region typically seen in the entorhinal cortex [9]. There was noticeable thinning over time in the anterior region of this subject’s rhinal cortex. Subject 2 started with a thickness profile similar to what was seen in the average stable NC. There was noticeable thinning over time in the lateral anterior region of the rhinal cortex, with particularly severe atrophy at the last scan time. Similarly, in Subject 3, we see a particularly thin posterior cortex and progressive thinning in the anterior lateral region of the rhinal cortex.

**Figure 5:**
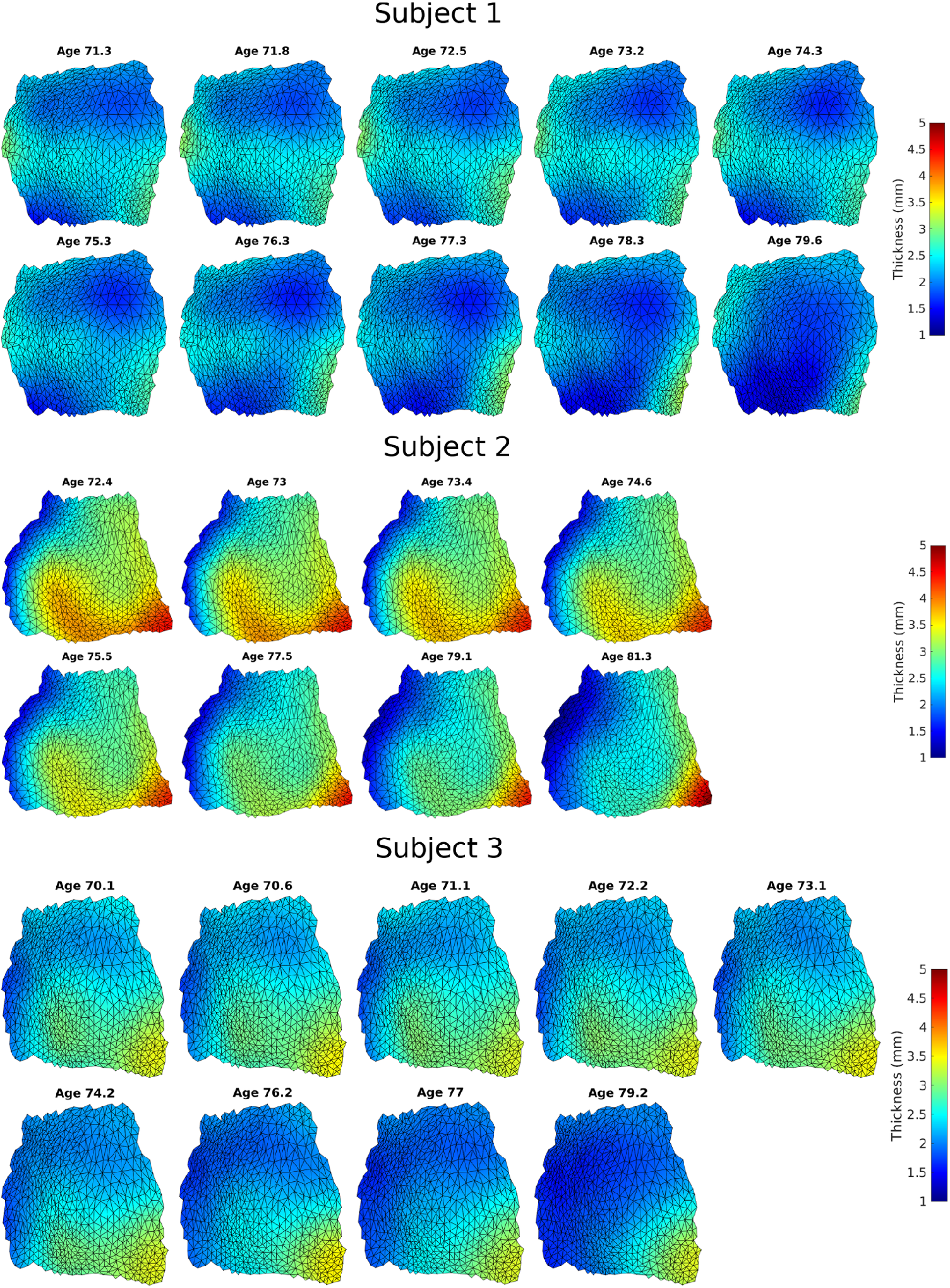
Cortical thickness over time of 3 subjects. Orientation of the surface is left (lateral), right (medial), top (posterior), bottom (anterior).

**Figure 6:**
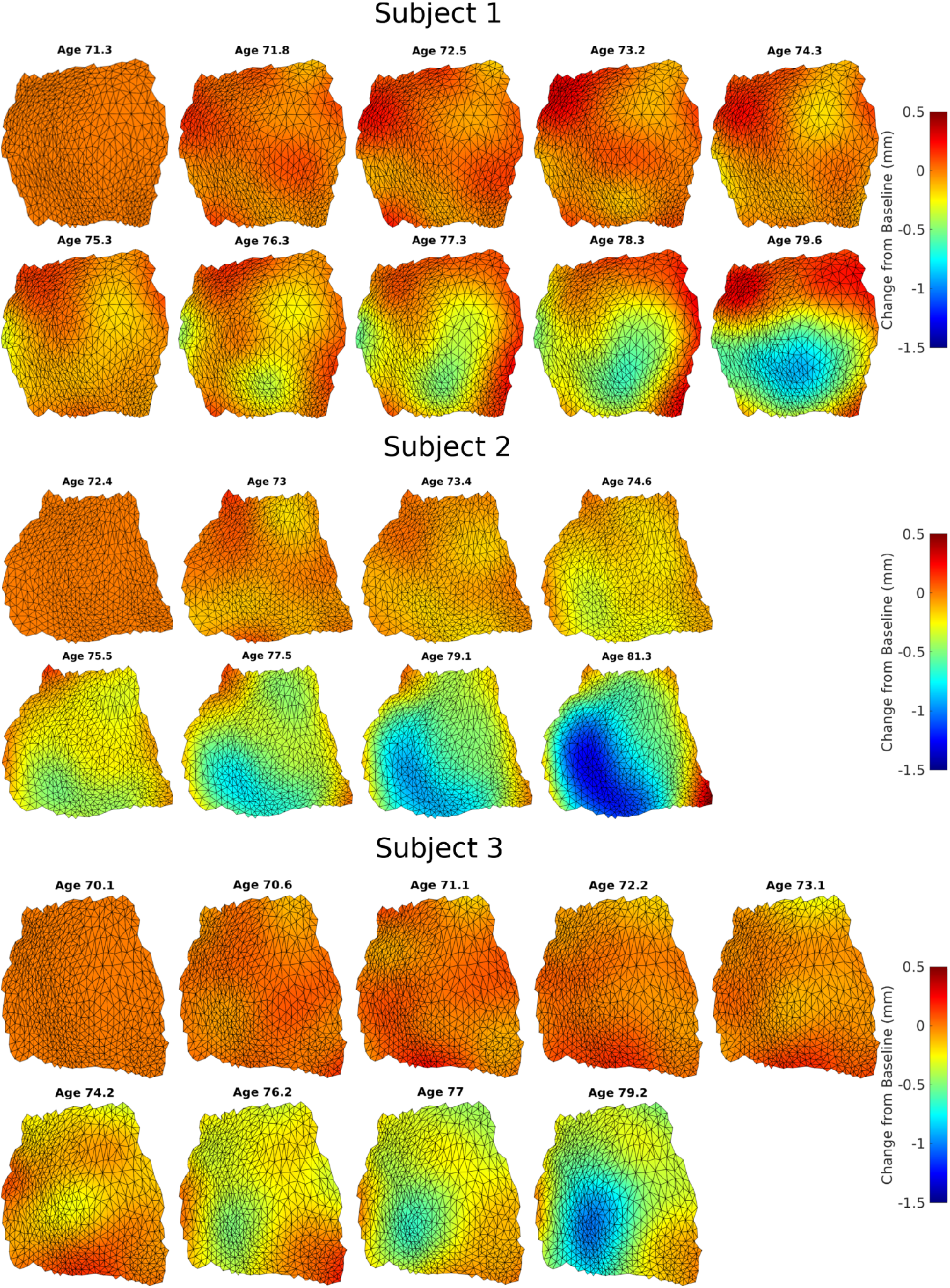
Change in cortical thickness from the first scan for 3 subjects. Orientation of the surface is left (lateral), right (medial), top (posterior), bottom (anterior).

For each subject, we estimated variation in cortical thickness over the rhinal cortex, also called shape difference, based on thickness measures from their first scan. The 3 subjects were individuals who did not convert to MCI within an 8 year period from their first scan. This is important because disease-related atrophy is thought to begin 8 to 10 years prior to symptom onset. The shape of the cortex at the first scan is a reasonable approximation of the cortical thickness prior to disease-related atrophy for these subjects [24], [11].

Parameters were estimated for 100 runs initialized from random guesses in the parameter space. The MAP estimator for each subject is shown in Table 2. All three subjects had an atrophy origin, *c*_0_, in the anterior lateral quadrant of the rhinal cortex. The speed was relatively consistent between subjects, varying from 0.41 mm/year and 0.81 mm/year. The diesease intensity also seemed to be relatively consistent between subjects, varying from 0.26 to 0.38. The slope and disease stage varied more widely, as did the initial average cortical thickness, 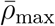. This variation in cortical thickness is expected, as seen in the cortical thicknesses of subjects with NC.

**Table 2:** MAP estimate of parameters for Subjects 1, 2, and 3.

| Parameter | Subject 1 | Subject 2 | Subject 3 |
| --- | --- | --- | --- |
| speed (mm/year) | 0.41 | 0.81 | 0.77 |
| slope (1/mm) | 0.56 | 2.06 | 5.36 |
| disease intensity | 0.38 | 0.26 | 0.32 |
| atrophy origin | [-1.35, -2.29] | [-6.11, -1.58] | [-3.17, -0.49] |
| disease stage | -5.66 | -8.23 | -9.96 |
| thickness (mm) | 2.72 | 3.66 | 3.01 |

We also examined the disease stage at the final observation time for the three subjects. Specifically, we examined the maximum disease stage across the rhinal cortex, which was consistenly in the anterior lateral region. The disease stage was −0.74 for Subject 1, −0.26 for Subject 2, and −0.64 for Subject 3. This corresponds to 32.3%, 43.5% and 34.5% disease-related atrophy at the time of the last scan.

Table 3 shows the 90% confidence intervals for these same parameters. The radius that circumbscribed 90% of atrophy origin, *c*_0_, was 0.61 mm for Subject 1, 0.59 mm for Subject 2, and 0.61 mm for Subject 3. The radius for atrophy origin and the confidence interval for speed, disease intensity, and average cortical thickness were relatively narrow in all three subjects. Slope and initial disease stage had somewhat larger confidence intervals. These findings are consistent with the results seen when estimating confidence intervals on the disease simulation.

**Table 3:** 90% confidence interval for Subjects 1, 2, and 3.

| Parameter | Subject 1 | Subject 2 | Subject 3 |
| --- | --- | --- | --- |
| speed (mm/year) | (0.38, 0.47) | (0.78, 0.95) | (0.67, 0.97) |
| slope (1/mm) | (0.47, 0.65) | (1.56, 2.76) | (4.09, 6.34) |
| disease intensity | (0.30, 0.43) | (0.25, 0.27) | (0.31, 0.32) |
| disease stage | (-5.36, -5.88) | (-7.82, -8.62) | (-9.52, -9.99) |
| thickness (mm) | (2.70, 2.75) | (3.64, 3.69) | (2.99, 3.03) |

## 5. Discussion

In this work, we have introduced a biologically-inspired subject-specific model of atrophy spread for Alzheimer’s disease. This is the first study, as far as we know, that examines the speed of atrophy spread. For the three subject case studies, the speed ranged from 0.4 mm per year to 0.8 mm per year, suggesting a relatively slow progression of atrophy across the rhinal cortex.

The origin of atrophy was also consistently found in the anterior lateral region of the rhinal cortex. This is in line with our previous population-wise studies that showed atrophy of preclinical subjects in the lateral region of the rhinal cortex, and more atrophy in the anterior region than the posterior region [11].

The initial disease stage is difficult to estimate accurately when the observation window is decades after the time of disease initialization. However, disease stage at the time of observations may prove to be more accurate. The disease stage at the final scan time in the three case studies suggest more than 32% atrophy occurred in the anterior lateral region of the rhinal cortex prior to a diagnosis of MCI. This is more atrophy than our population study of preclinical subjects showed, where the disease model had a maximum atrophy of 17% at the time of MCI diagnosis [11]. One possible explanation for this discrepancy may be that population-wise studies attribute some disease-related atrophy to individual variability in cortical thickness, since no prior is available on an individual basis of differences in thickness. On the other hand, subject-specific modeling with subjects that are scanned over a longer period of time benefit from a personalized model of cortical thickness over the rhinal cortex.

This study has several strengths. The proposed mechanistic model was biologically-inspired by properties of neurofibrillary tau tangles and localization of this protein as seen in autopsy studies of Alzheimer’s disease. Tests conducted on simulated data suggest we can obtain reasonable accuracy given a window of 5 to 10 years of observations. The three case studies were selected carefully to have a long duration of observations and strong evidence of early Alzheimer’s disease.

However, the study does have a few limitations. The model assumed uniform spread of atrophy across the surface. It is not known whether atrophy spreads uniformly or whether there is some bias, for example, for atrophy to move faster lateral to medial than anterior to posterior.

The cortical thickness profile prior to disease was estimated for each subject based on their first scan. This strategy cannot be employed on subjects with a shorter duration of observation, where it is not known whether disease-related atrophy has occurred. Developing a subject-specific prior of cortical thickness is a topic of interest since collateral sulcus location and cortical thickness vary greatly from subject to subject.

As a greater amount of high resolution T2 data is made available, it may also be possible to extend this analysis to the subiculum and subregions of the hippocampus, such as CA1.

In summary, we have introduced a new model of atrophy spread that can be used to estimate disease-related parameters such as the original of disease atrophy and speed of spread for an individual.

## 6. Materials and Methods

### 6.1. Data processing

15 subjects with stable normal cognition (NC) were selected from the ADNI database (adni.loni.use.edu) for estimation of cortex shape and age-related atrophy. The criteria for NC subjects was a clinical dementia rating equal to 0 on all annual evaluations, evidence of performance within the normal range on the Logical Memory Subtest of the Wechsler Memory Scale on all annual evaluations, and a negative result for elevated amyloid *β* levels on the baseline evaluation (as established by the ADNI Biospecimen Core). All subjects were scanned on 3T MRI for 2 or more years.

3 subjects were selected from the ADNI database for case studies of the atrophy spread model. We screened for subjects that met the following criteria:

- labeled a control at baseline
- remained a control for at least 8 years
- were scanned on 3T MRI for at least 5 years
- showed evidence of risk for MCI conversion

– converted to MCI after 8 or more years
– subjective memory complaint and amyloid positive
– subjective memory complaint and impaired delayed recall when no amyloid data available

Table 4 shows the age, sex, and scan information of the subjects in this study. Scans obtained using a new accelerated protocol in ADNI 3 were excluded from analysis to mitigate concerns of differences in cortical thickness by scan protocol. Subject 1 was followed for 12 years and converted to mild cognitive impairment at year 9 or 10 (no evaluation was performed at year 9). Subject 2 was followed for 9 years and did not convert to MCI during this period of observation. The subject was amyloid positive at baseline (as established by the ADNI Biospecimen Core) with an increasing subjective memory complaint (as measured by the Everyday Cognition Questionnaire). Subject 3 was followed for 11 years and did not convert to MCI during this period of observation. The subject had no CSF sample analyzed, so amyloid data was not be obtained. The subject had an increasing subjective memory complaint (as measured by the Everday Cognition Questionnaire) and a delayed recall score below the normal range on both the Logical Memory Subtest of the Wechsler Memory Scale (with a score as low as 7), and the Rey Auditory Verbal Learning Test (with a score as low as 2).

**Table 4:** Demographic data (mean ± standard). *converted to MCI at year 9 or 10. Note that the scan period is often shorter than the follow-up period because accelerated 3T scans introduced in ADNI3 were excluded from analysis.

|  | Stable NC | Subject 1* | Subject 2 | Subject 3 |
| --- | --- | --- | --- | --- |
| Number of Subjects | 15 |  |  |  |
| Age (years) | 71.44 ± 7.17 | 71.3 | 72.4 | 70.1 |
| Number of Scans | 4.53 ± 0.52 | 10 | 8 | 9 |
| Scan Period (years) | 3.36 ± 1.14 | 8.3 | 8.9 | 9.1 |
| Follow-up Period (years) | 5.67 ± 3.22 | 12.1 | 8.9 | 10.6 |
| Sex (% Female) | 46.67% | F | F | M |

To calculate cortical thickness, we used previously established methods [11] and briefly summarize the methods here. We followed a landmark-based protocol for manual segmentation of the left ERC and TEC. A template mesh for the shape of the ERC plus TEC surface was generated from a set of segmentations of control and preclinical subjects by calculating the average diffeomorphism in a Bayesian setting. For each subject, we adjusted for variability in segmentation boundaries across scans using an approach called unbiased longitudinal diffeomorphometry [20]. An online tool to perform these steps are available on https://www.mricloud.org.

Figure 7 shows an example mapping of the population average template surface (in gray) to a subject’s set of data (in red) using unbiased longitudinal diffeomorphometry. At each iteration, the geodesic mapping from the population template to subject template, the insertion time t*, and the geodesic mapping through all the timepoints from t* is updated. The result is a set of surfaces with corresponding vertices smoothed for variation over time.

**Figure 7:**
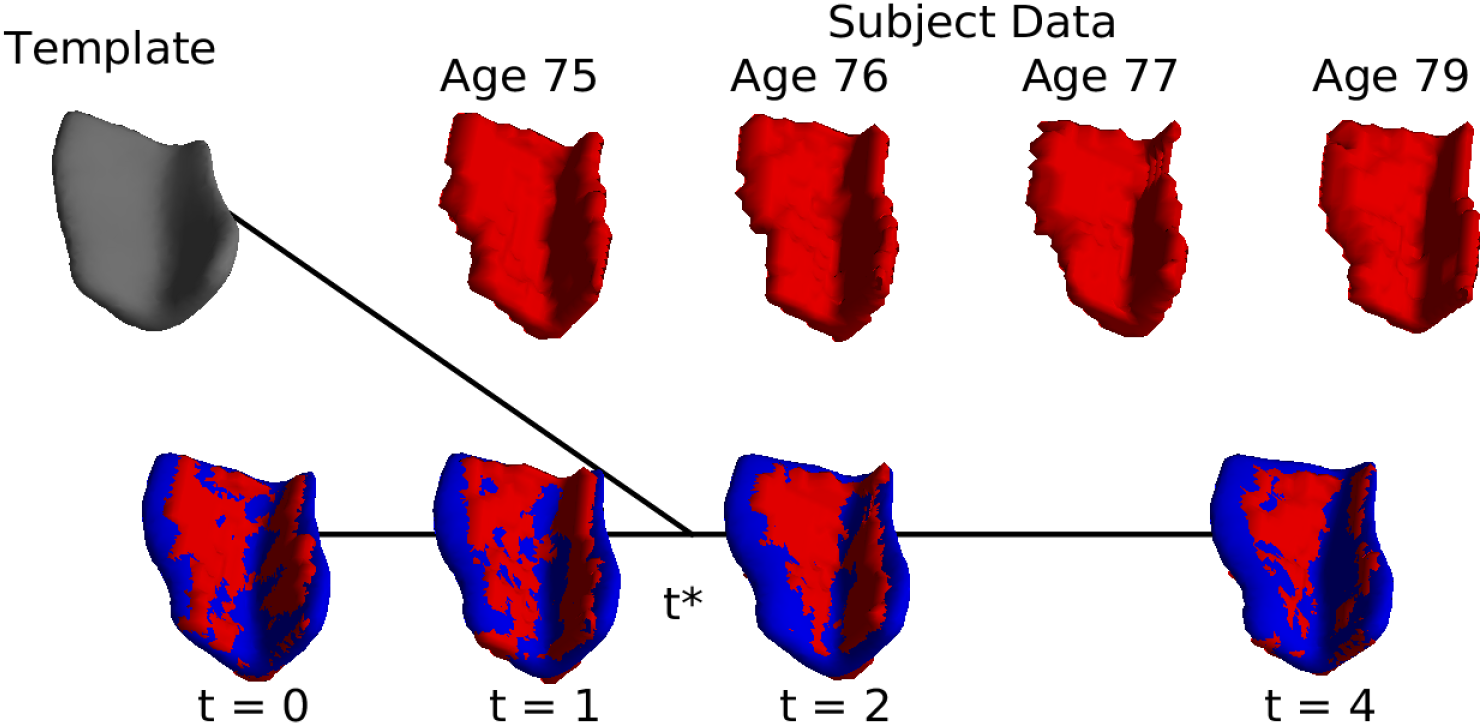
unbiased longitudinal diffeomorphometry

The triangulated meshes were then cut using a semi-automated approach into two surfaces: the gray matter-white matter boundary surface and the pial surface (gray matter-cerebral spinal fluid boundary surface). We calculated the cortical thickness from one surface to the other using an approach called normal geodesic flow [18]. This results in a set of vertices with a measure of cortical thickness, where the vertices have one-to-one correspondence over all scans and all subjects. We further reduced the dimensions of the vertices from three to two using multi-dimensional scaling. The code used to perform cortical thickness calculation is available for download at https://www.bitbucket.org/laurent_younes/registration/src/master/py-lddmm/.

Figure 8 shows an example of how cortical thickness was calculated on the population average template surface. First, the surface was cut into the CSF-GM boundary (inferior) and GM-WM boundary (superior) surfaces. We calculate the cortical thickness for each vertex on the CSF-GM boundary surface using normal geodesic flow. Cortical thickness is the distance traveled along the geodesic flow, as shown in black lines in Figure 8b. The vertex-wise measures of cortical thickness are shown in color on the flattened CSF-GM boundary surface in Figure 8 c. Figures of flattened surfaces in the following section are oriented as shown here in Figure 8 c.

**Figure 8:**
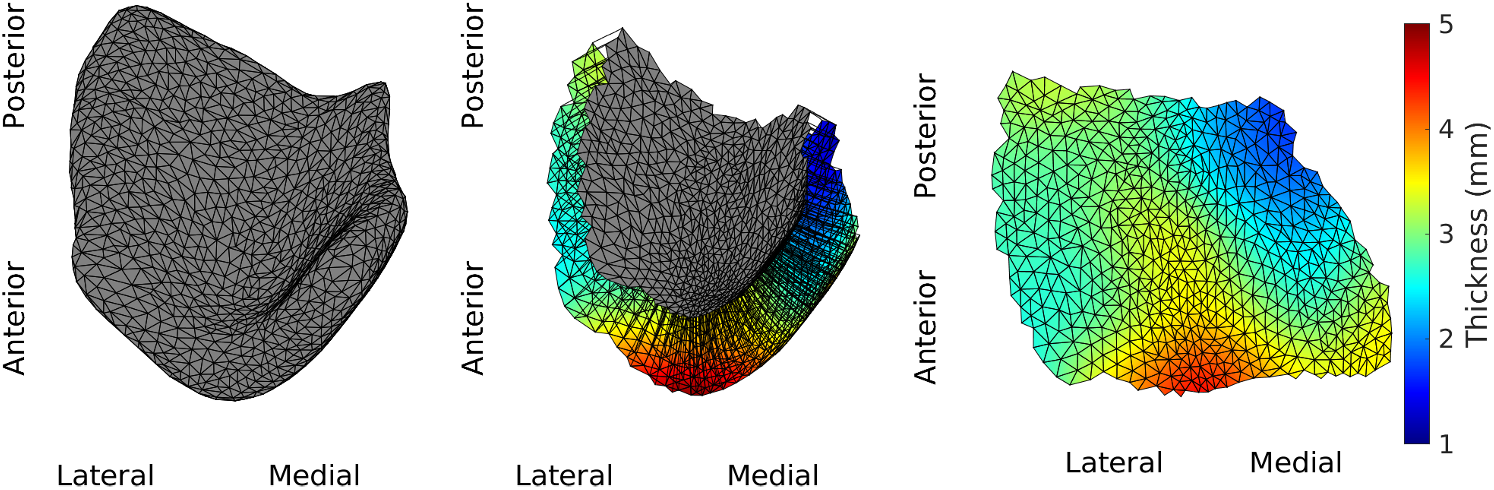
Population-based surface template of ERC plus TEC (left), template cut with normal geodesic flow trajectories (middle), flattened template with cortical thickness measures in color (right)

### 6.2. Disease simulation generation

The shape of the cortex *h* in Eqn 4 and age-related atrophy rate λ_0_ were estimated a priori. A mixed effects model was used to estimate the cortex shape (variation in thickness over vertices), the fixed effects of age and sex, and the variance of zero-mean Gaussian noise *ϵ*, based on 15 stable NC subjects. The model of cortical thickness (thk) for a subject *i*, scan *j*, at vertex *k* is shown in Eqn 5. The parameter for cortical thickness 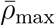 was set to be the average cortical thickness from this model, 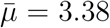. The constant for age-related atrophy was set to the vertex-wise solution λ_0_ from this model.

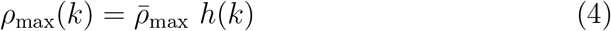

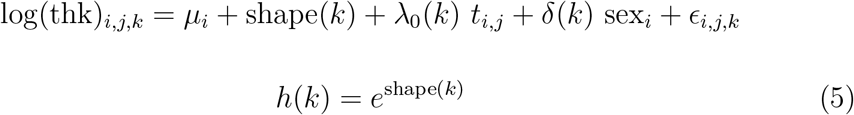

We simulated disease-related atrophy by generating cortical thickness over time using the discretized forward model in Eqn 9 on the population template surface. Disease-related parameters *c*_0_, *b*_0_, λ_1_, *κ*, and *D* were chosen to reflect characteristics of Alzheimer’s disease as best as is currently understood.

The initial seeding of local activity *c*_0_ was chosen to be in the anterior lateral region, where studies have shown neurofibrillary tangles first form [3], and where early disease-related atrophy was observed from MRI data in our previous studies [10], [11]. The initial disease stage *b*_0_ was set to –5 such that disease-related atrophy, 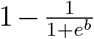, was less than 1% when the model initializes at age 50. Note that all subjects in this study were enrolled asymptomatic starting at age 65 and older; no disease-related atrophy is expected at age 50 in these cases.

To select a reasonable estimate of disease intensity λ_1_, we considered evidence from a set of 19 subjects diagnosed with dementia, the latest clinical stage in the disease course. Compared against a set of 33 NC subjects, these subjects showed evidence of 15.1% atrophy throughout the ERC plus TEC at the time of diagnosis and a rate of 4.05% disease-related atrophy per year. Next, we considered a simplified model where b is constant across the surface, the time step *dt* = 1 year. We approximated λ_1_ = 0.09 in the following way:

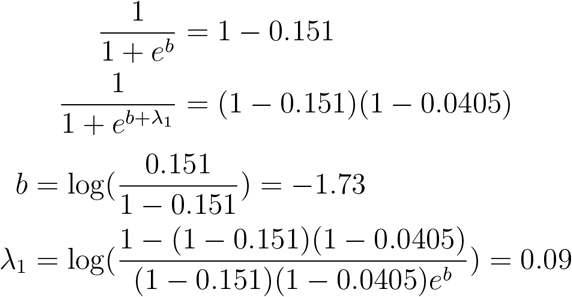

There is little data to related to the selection of *κ* and *D*. We selected *κ* = 1 and *D* = 1. Note that 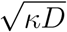 is the speed of spread across the surface, and 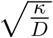 is the slope of local activity rate *a* at the wave front. *κ* and *D* were chosen such that local activity rate *a* traverses the surface in 10 years with a slope of 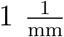. The final parameters chosen to construct the disease simulation were as follows:

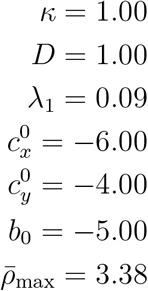

The information in this section was also used to inform boundaries for randomized initial guesses of parameters. The initial guess for *c*^0^ was selected such that 95% of the volume of Gaussian *a*_0_ centered about *c*^0^ is on the surface. This excludes points too close to the surface boundary. The initial guesses for other parameters were selected from the following ranges:

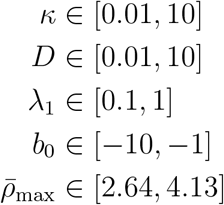

### 6.3. Parameter estimation approach

To reduce the number of parameters estimated to a reasonable size, we constrain the initial conditions and cortex shape. The initial local activity rate *a*^0^ is constrained to be Gaussian with a center at *c*^0^ and fixed variance *σ*^2^ = 1, as shown in Eqn 6.

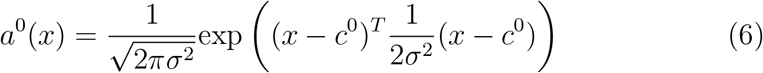

The initial disease stage 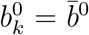, a constant across all *k*. The smaller 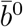 is, the more resistant a subject is to cortical thinning early on as the disease progresses. The model is also fixed to begin at age 50. The simplified model parameters *θ* are shown in Equation 7.

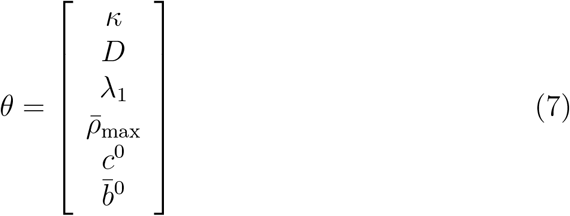

We introduce new notation in Eqn 8, to re-write the atrophy spread model as Eqn 9.

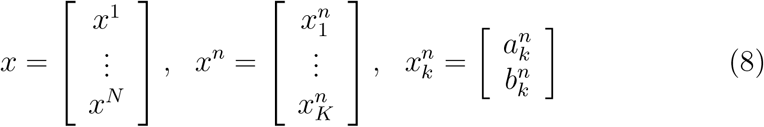

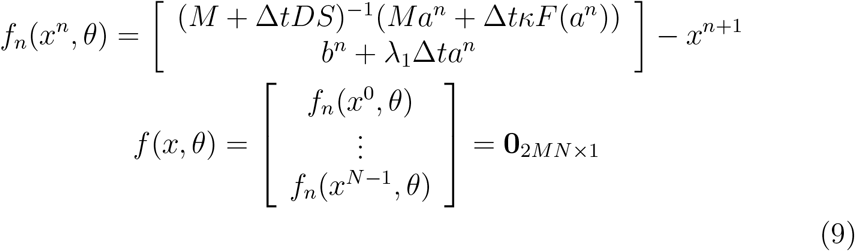

We solve for *θ* that minimizes *g*(*x, θ*), the squared difference between observed and modeled cortical thickness. This is a constrained optimization problem with constraints *f*(*x, θ*) = 0 and cost function *g*(*x, θ*). We reformulate and solve the problem in its dual form using the adjoint state method. Details on this method and its derivation are in the Supplementary Materials 7.2. The final set of equations used to perform gradient descent and estimate parameters, *θ*, are shown in Eqn 10.

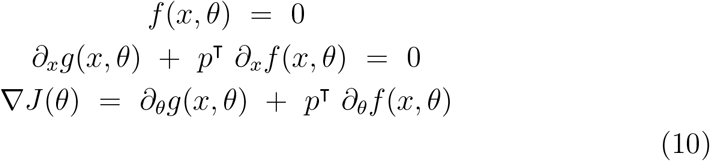

### 6.4. Analysis on posterior distribution

The goal of this analysis is to build a 90% confidence interval for each model parameter. In order to calculate these confidence intervals, we want an explicit expression of the posterior distribution, *p*(*θ* | *ρ*_observed_). We begin with the probability of observing *ρ*_observed_ given a set of parameters, as shown in Eqn 11. Note that *ρ*(*θ*) is shorthand for the model of atrophy spread with input parameters *θ* and output cortical thickness *ρ*.

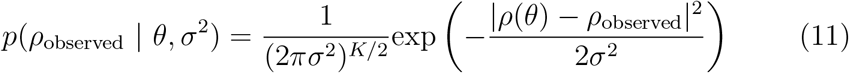

We have introduced a nuisance parameter, *σ*^2^, which represents the noise associated with observations. We can approximate the distribution of *σ*^2^ as an inverse Wishart distribution with mean and variance 1, as shown in Eqn 12.

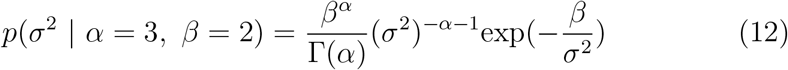

The values for *β* and *α* associated with an inverse Wishart distribution with mean and variance equal to 1 were derived in the following way:

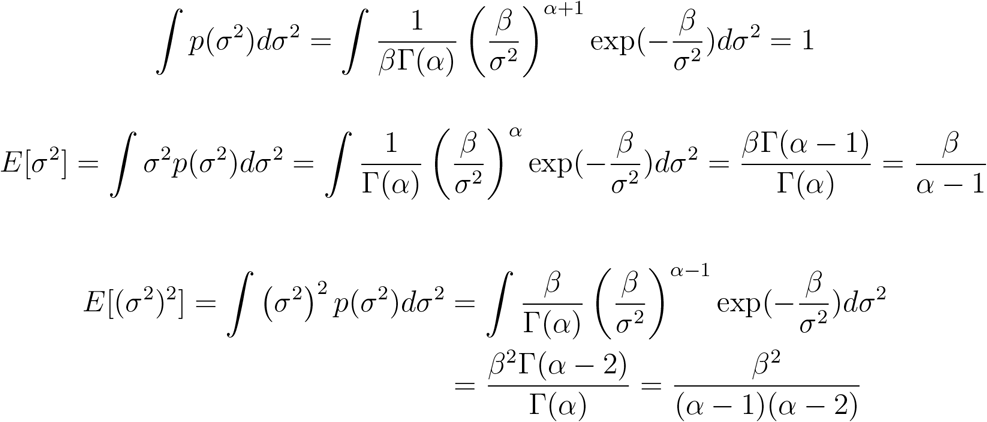

Setting the mean to 1, we see *β* = *α* – 1. Then, setting the variance to 1 and substituting *β* = *α* – 1, we see that *α* = 3, *β* = 2.

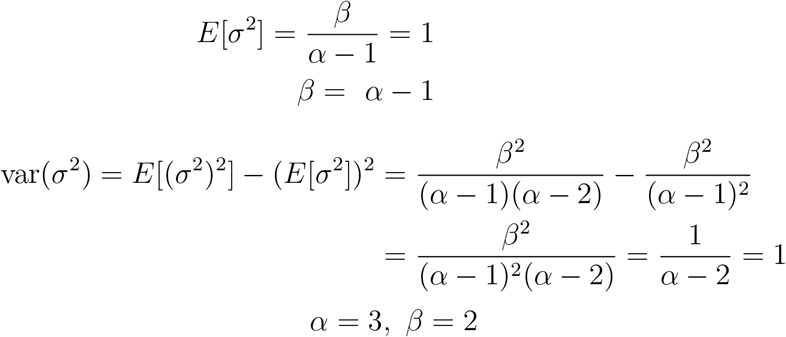

This approximation allows us to integrate for an explicit expression of a value proportional to the prior distribution *p*(*θ* | *ρ*_observed_) over *K* vertices, as derived below. *p*(*θ*) is uniform across the parameter space. The final expression proportional to the approximate posterior distribution with respect to *K, α*, and *β* is shown in Eqn 13.

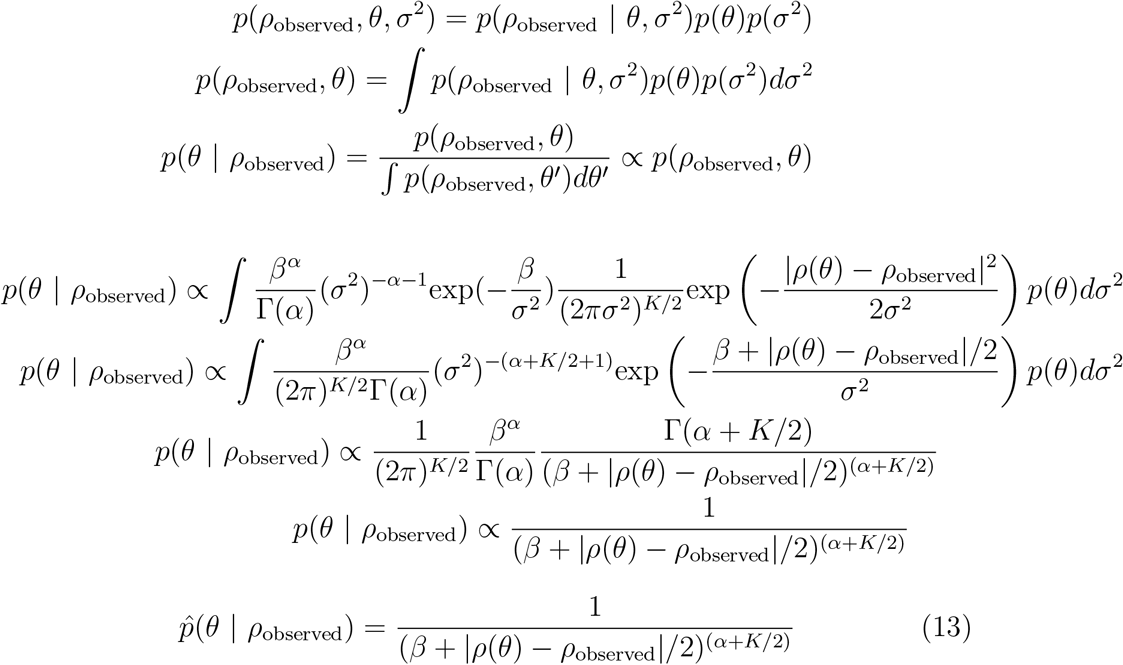

Directly sampling the posterior distribution was too computationally intensive for this high dimensional parameter space. Instead, we implemented a Markov chain Monte Carlo algorithm, called the Metropolis-Hastings algorithm, to randomly sample from the posterior distribution. We start from a random initial guess *θ*_0_ in the parameter space. For each of 20, 000 iterations, we select a candidate 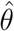 by updating the current *θ* with Gaussian noise of fixed variance. The variances were selected empirically to be: 0.01 for *k*, 0.01 for *D*, 0.001 for λ_1_, 0.01 for *c*_0_ in the x and y direction, .05 for 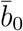, and .005 for 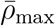. As shown in Eqn 14, we update *θ* to 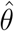 with a probability that is the ratio of the prior distribution at *θ* and 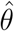, given *α* = 3, *β* = 2, and observations *ρ*_observed_ for *K* vertices.

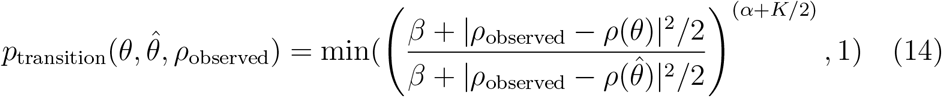

The 20,000 samples of *θ* are then used to calculate the 90% confidence intervals, and the radius of a circle that circumscribes 90% of *c*_0_ values.

## 7. Supplementary Material

### 7.1. Discretization of atrophy spread model

To derive the discretized model, we start with a variational formulation of Eqn 3, as shown in Eqn 15. We introduce a new piecewise continuous function *ϕ*(*s*) ∈ *H*^1^ (Ω), where *H*^1^ is a Hilbert space.

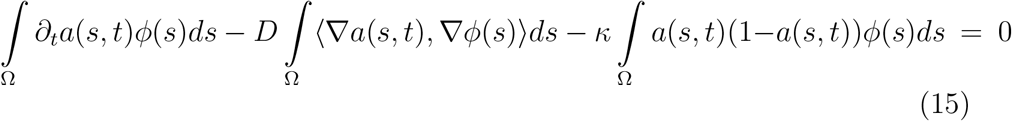

Let *k* ∈ 0, 1,…, *K* represent indices of the discretized space, more specifically a triangulated mesh. We can represent local activity rate *a*(*s, t*) as a sum of spatially discretized rates *a_k_*(*t*) weighted by function *φ_k_*(*s*) as shown in Eqn 16. *ϕ_k_*(*s*) is a tent function with a constant gradient ∇*ϕ_k_*(*s*) = ∇*ϕ_k_* for all s within a triangular face of the mesh.

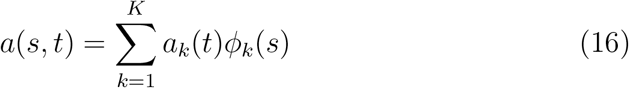

Using Eqn 15 and Eqn 16, we can now represent this system in matrix form with mass matrix *M*, stiffness matrix *S*, and nonlinear reaction term *F* as shown in Eqn 17.

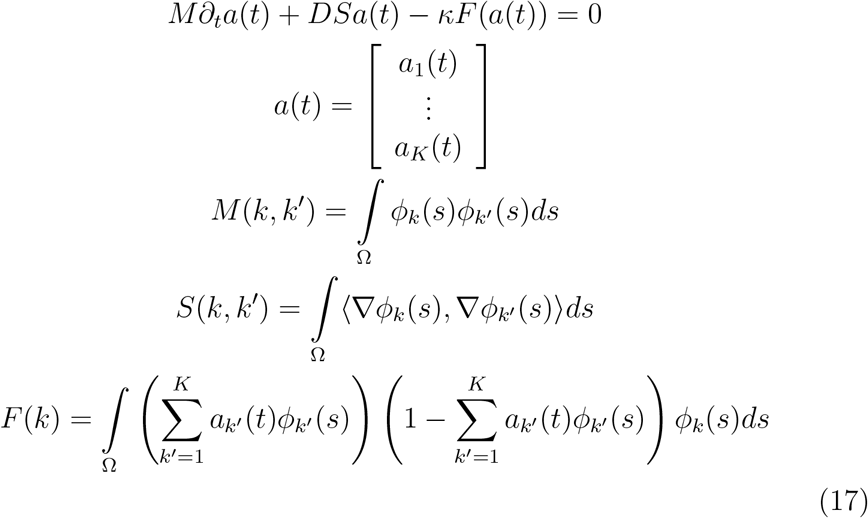

Computing *M, S*, and *F* as shown above is computationally intensive. Instead, we calculate an approximation using the properties of *ϕ*. Let *R* = {*r_u_, r_v_, r_w_*} be the set of 2-D vertices of a triangle. Let *r_uv_* be the midpoint of vertex *r_u_* and *r_v_*, and *r_uvw_* be the barycenter of the triangle. The area of the triangle is then Eqn 18.

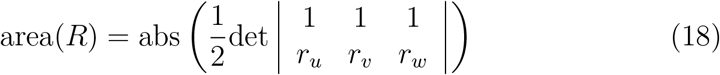

We now can introduce 1^st^, 2^nd^ and 3^rd^ order approximations of *ϕ*(*s*) over a triangle in Eqn 19, Eqn 20, and Eqn 21 respectively.

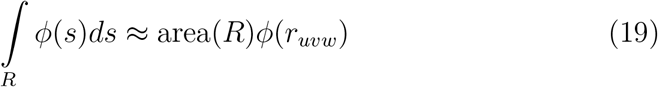

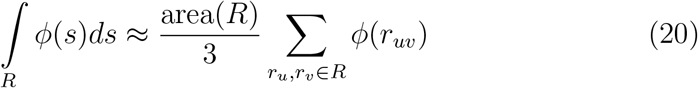

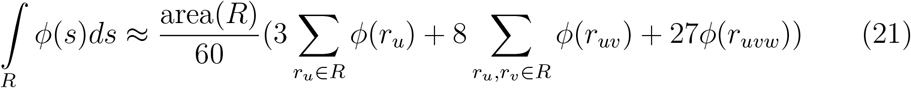

*ϕ* is a 1^st^ order piecewise linear function. Specifically, note that *ϕ_k_* (*r_k_*) = 1, 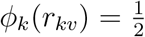 and 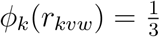. *M* can be represented as a 2^nd^ order function, *S* can be represented as a 1^st^ order function, and *F* can be represented as a 3^rd^ order function summed over all the triangles *R* as shown in Eqn 22, Eqn 23, and Eqn 24 respectively.

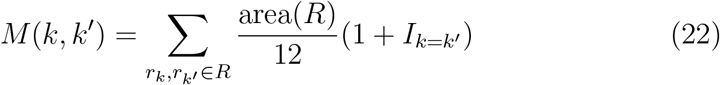

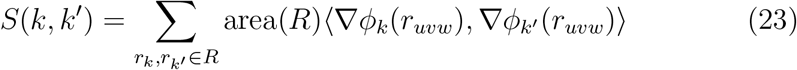

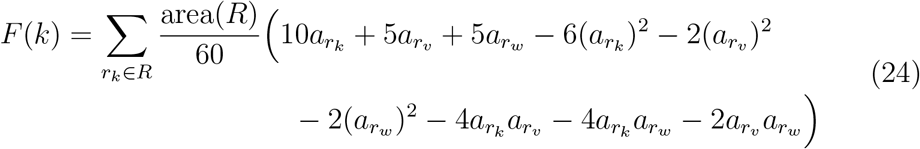

Let *n* ∈ 0,1,…, *N* represent indices of discretized time with a step size of Δ*t*, and let us introduce a simplified notation where *a^n^* = *a*(*t_n_*). We discretize Eqn 17 using a semi-implicit scheme, as shown in Eqn 25.

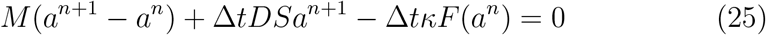

The final set of discretized equations are shown in Eqn 26.

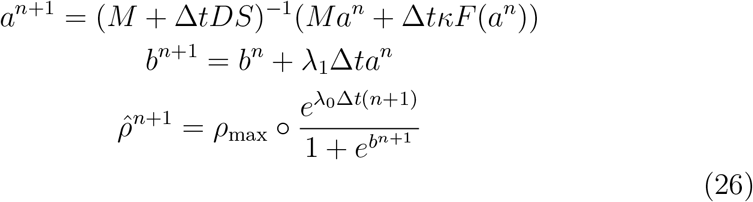

### 7.2. Derivation of parameter estimation approach

We reformulate and solve the constrained optimization problem in its dual form, using the adjoint state method. The derivation of this new formulation is shown below.

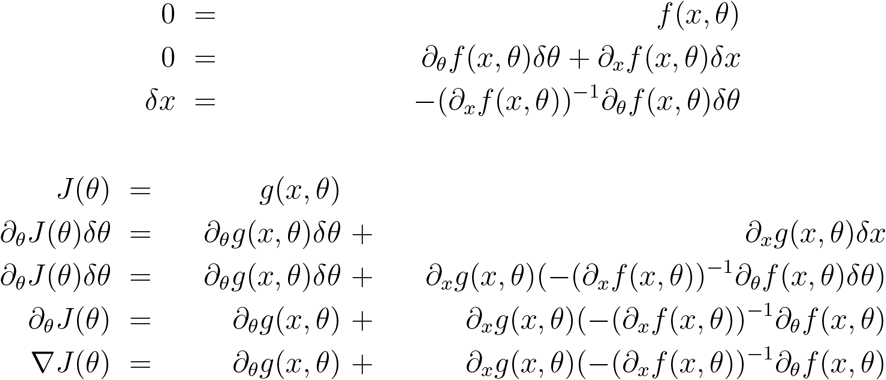

Next we introduce the adjoint state vector, *p*, as shown in Eqn 27.

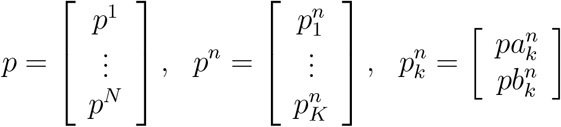

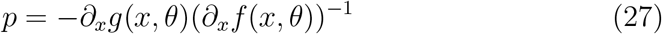

The set of equations for parameter estimation are shown in Eqn 28. The general implementation is as follows: Given an initial guess *θ*, calculate *x* recursively using the constraint in Line 1 of Eqn 28. Next, calculate the adjoint state vector *p* recursively using Line 2 of Eqn 28. The explicit recursive relationship to solve for *p* is Eqn 29. Finally, calculate the gradient ∇*J*(*θ*). Update *θ* iteratively using gradient descent.

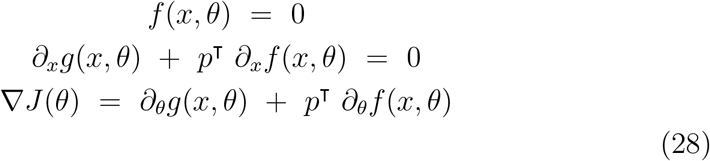

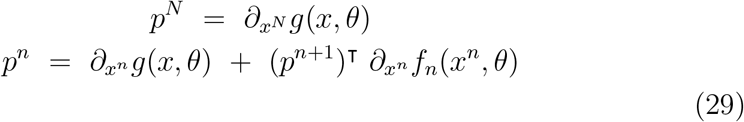

The solution for ∇*J* in terms of variables *a, b, p, θ*; matrices *M, S, F, dF*; and constants λ_0_, Δ*t, h* is shown in Equation (30).

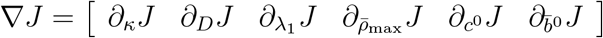

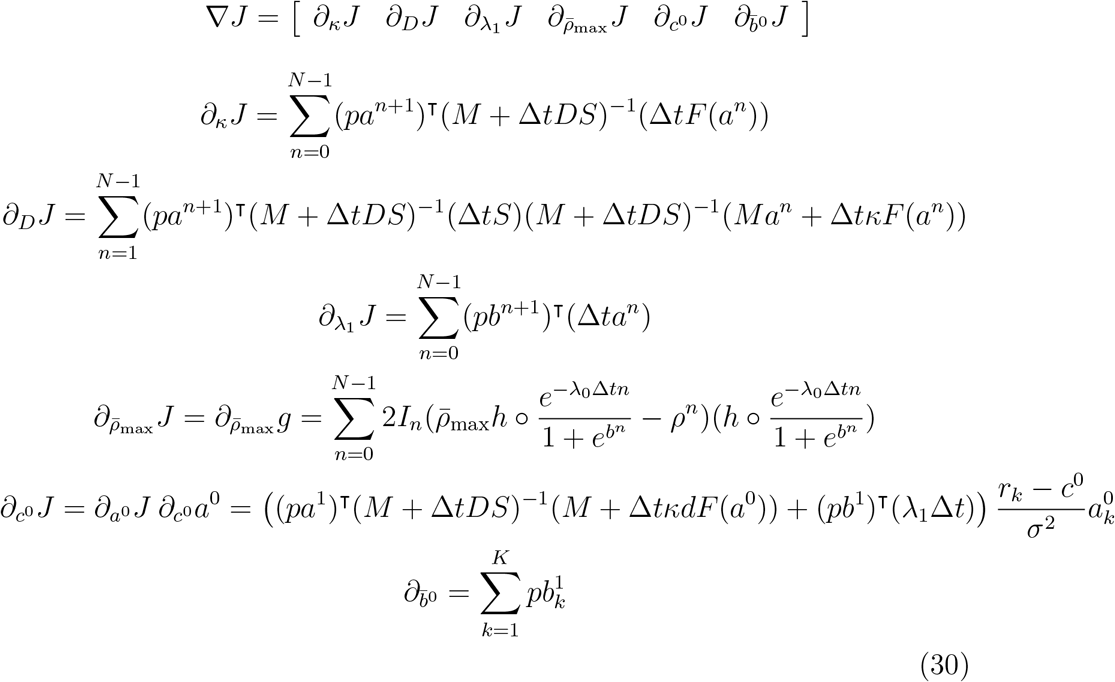

where *r_k_* = [*r_k,x_ r_k,y_*] location of vertex *k*

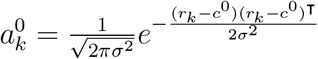

Model implementation is available for download from https://www.github.com/sue-kulason/FKPP.

## 8. Acknowledgments

We thank Arnold Bakker for his invaluable contributions in subject selection and Eileen Xu for her careful segmentions of the entorhinal and transentorhinal cortex prepared in this study.

## 9. Funding and Support

This work was supported by the National Institutes of Health [grant number P41 EB015909]; the National Institute on Aging [grant number R01 AG048349]; the Kavli Neuroscience Discovery Institute; and Fulbright France. Michael Miller reports personal fees from AnatomyWorks, LLC, outside the submitted work, and jointly owns AnatomyWorks. This arrangement is being managed by the Johns Hopkins University in accordance with its conflict of interest policies. Dr. Miller’s relationship with AnatomyWorks is being handled under full disclosure by the Johns Hopkins University.

Data used in the preparation of this article were obtained from the Alzheimer’s Disease Neuroimaging Initiative (ADNI) database (adni.loni.usc.edu). As such, the investigators within the ADNI contributed to the design and implementation of ADNI and/or provided data but did not participate in the analysis or writing of this report. A complete listing of ADNI investigators can be found at http://adni.loni.usc.edu/wp-content/uploads/how_to_apply/ADNLAcknowledgement_List.pdf

Data collection and sharing for this project was funded by the Alzheimer’s Disease Neuroimaging Initiative (ADNI) (National Institutes of Health Grant U01 AG024904) and DODADNI (Department of Defense award number W81XWH-12-2-0012). ADNI is funded by the National Institute on Aging, the National Institute of Biomedical Imaging and Bioengineering, and through generous contributions from the following: AbbVie, Alzheimer’s Association; Alzheimer’s Drug Discovery Foundation; Araclon Biotech; BioClinica, Inc; Biogen; Bristol-Myers Squibb Company; CereSpir, Inc; Cogstate; Eisai Inc; Elan Pharmaceuticals, Inc; Eli Lilly and Company; EuroImmun; F. Hoffmann-La Roche Ltd and its affiliated company Genentech, Inc; Fujirebio; GE Healthcare; IXICO Ltd; Janssen Alzheimer Immunotherapy Research & Development, LLC; Johnson & Johnson Pharmaceutical Research & Development LLC; Lumosity; Lundbeck; Merck & Co, Inc; Meso Scale Diagnostics, LLC; NeuroRx Research; Neurotrack Technologies; Novartis Pharmaceuticals Corporation; Pfizer Inc; Piramal Imaging; Servier; Takeda Pharmaceutical Company; and Transition Therapeutics. The Canadian Institutes of Health Research is providing funds to support ADNI clinical sites in Canada. Private sector contributions are facilitated by the Foundation for the National Institutes of Health (www.fnih.org). The grantee organization is the Northern California Institute for Research and Education, and the study is coordinated by the Alzheimer’s Therapeutic Research Institute at the University of Southern California. ADNI data are disseminated by the Laboratory for Neuroimaging at the University of Southern California.

